# A computational model of the escape response latency in the Giant Fiber System of *Drosophila melanogaster*

**DOI:** 10.1101/435362

**Authors:** Hrvoje Augustin, Asaph Zylbertal, Linda Partridge

## Abstract

The Giant Fiber System (GFS) is a multi-component neuronal pathway mediating rapid escape response in the adult fruit-fly *Drosophila melanogaster*, usually in the face of a threatening visual stimulus. Two branches of the circuit promote the response by stimulating an escape jump followed by flight initiation. Our recent work demonstrated an age-associated decline in the speed of signal propagation through the circuit, measured as the stimulus-to-muscle depolarization response latency. The decline is likely due to the diminishing number of interneuronal gap junctions in the GFS of ageing flies. In this work, we presented a realistic conductance-based, computational model of the GFS that recapitulates our experimental results and identifies some of the critical anatomical and physiological components governing the circuit’s response latency. According to our model, anatomical properties of the GFS neurons have a stronger impact on the transmission than neuronal membrane conductance densities. The model provides testable predictions for the effect of experimental interventions on the circuit’s performance in young and ageing flies.

## INTRODUCTION

Escape responses are evolutionarily ancient mechanisms used by many species as their main defence against predator attacks. Intense selection pressure has led to dedicated reflex circuits that continuously monitor the environment for danger and trigger escape behaviours when presented with a specific set of threatening stimuli. These circuits must be able to respond within a minimal time frame to prevent capture and maximize chances of survival (Herberholz et al., 2004; Walker et al., 2005). Escape circuits are therefore characterized by extremely fast reaction times, with response latencies as short as a few milliseconds (Card and Dickinson, 2008; Dill, 1974). In dipteran insects, escape responses are mediated by the Giant Fiber System (GFS). Prompted by a visual (and, possibly, mechano-sensory) stimulus, the adult fruit-fly *Drosophila melanogaster* executes a stereotyped sequence of events that results in an escape jump followed by flight initiation (Allen et al., 2006; Fayyazuddin et al., 2006; Trimarchi and Schneiderman, 1995). The GFS consists of two descending, nonmyelinated Giant Fiber (GF) interneurons that originate in the brain, and downstream neurons that innervate and activate flight muscles (DLMs) and jump muscles (TTMs) (Allen et al., 2006; King and Wyman, 1980; Sun and Wyman, 1997) (Fig. 1). Functionally, electrical synapses are a dominant type of synapse in the *Drosophila* GFS (Phelan et al., 1996; Trimarchi and Murphey, 1997), with chemical (cholinergic) synapses playing a minor role (Allen and Murphey, 2007). Gap junctions are the physical substrate of electrical synapses that provide physical continuity between the cytoplasms of closely apposed pre- and post-synaptic neurons (Bennett, 1997). Compared to chemical synapses, transmission across electrical synapses is extraordinarily fast, with the possibility of the current flowing in either direction across the gap junction (Purves et al., 2001). Electrical synapses are therefore frequently found in places where fast transmission is critical, such as in escape response and motion-processing circuits (Cook and Becker, 1995). In the *Drosophila* GFS, the *shaking-B* gene (*shakB*, *inx8*) instructs the formation of heterotypic, unidirectional (rectifying) electrical synapses (Phelan et al., 1998; Stebbings et al., 2002; Wu et al., 2011).

**Figure 1.**
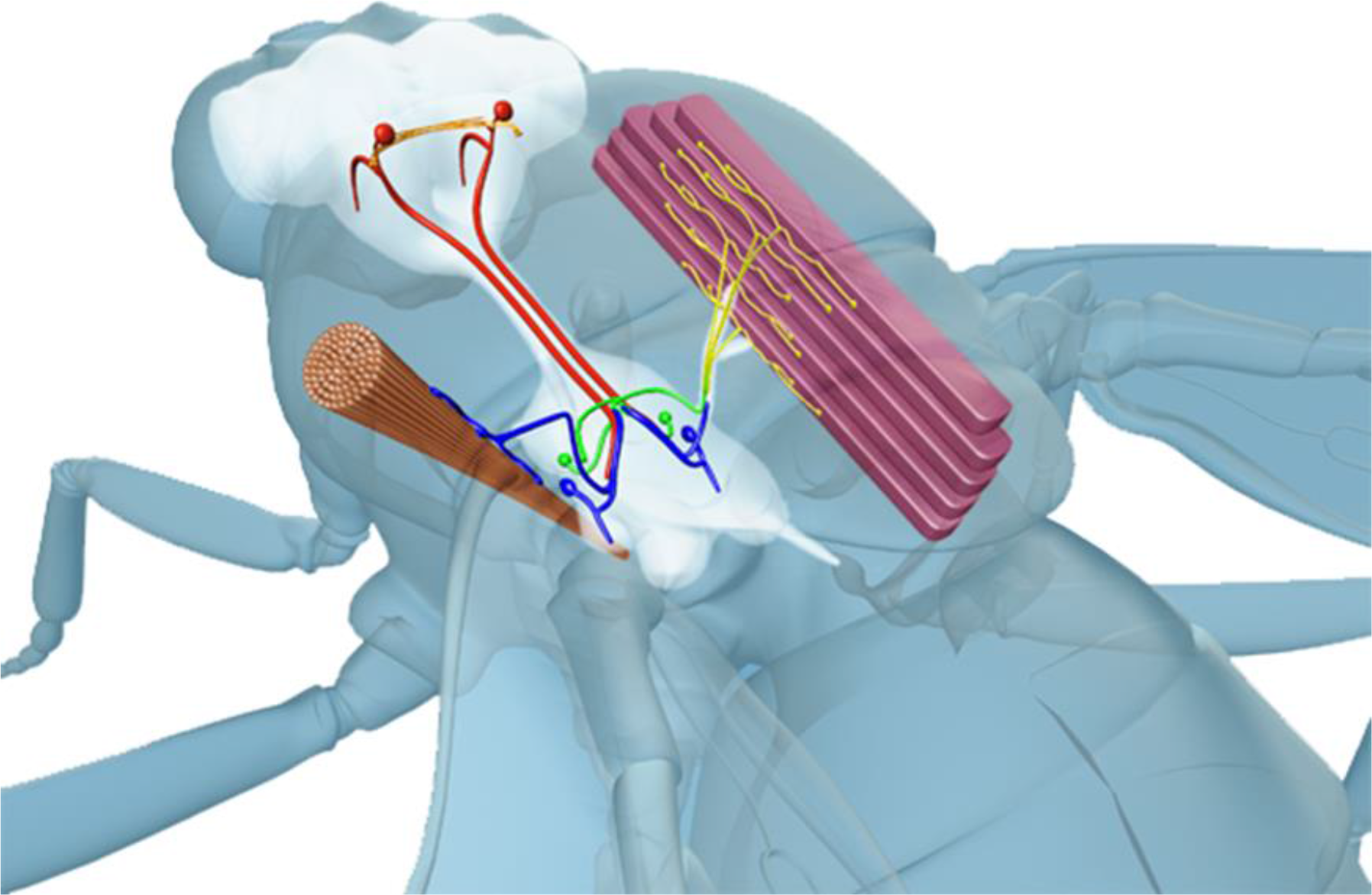
Diagram of the GFS anatomy. Two Giant Fiber (GF) interneurons originating in the brain (red) descend to the thoracic ganglia where they connect, via a mixed (electrical and chemical) synapse, to the tergotrochanteral motoneuron (TTMn, *blue*) innervating the cylindrical tergotrochanteral (jump) muscle (TTM). In the second branch of the circuit, the GFs form a mixed synapse with the peripherally synapsing interneuron (PSI, *green*) which, in turn, chemically synapses onto the dorsal longitudinal motoneurons (DLMns, *yellow*) innervating the dorso-longitudinal (flight) muscle (DLM). Red circles in the brain denote approximate positions of the GF cell bodies.

Loss of gap junctions in the nervous system occurs normally as a consequence of ageing. For example, astrocytic gap junctional plaques are drastically reduced in the brains of ageing mice (Cotrina et al., 2001), likely affecting inter-astrocytic and neuron-glia metabolic coupling (Cai et al., 2017). The structural proteins comprising the gap junctional channels are called connexins or pannexins in vertebrates (Hormuzdi et al., 2004) and innexins in invertebrate species (Hasegawa and Turnbull, 2014). In various knock-out mutants, widespread disruption of the neuronal gap junctional coupling leads to reduced synchronicity of neural networks (Deans et al., 2001), impaired oscillatory patterns in the brain (Buhl et al., 2003), neuronal hyperexcitability (Sutor et al., 2000), increased neuronal apoptosis (Nakase et al., 2003) and reduced neuroprotection after ischemic injury (Siushansian et al., 2001).

In our recent paper (Augustin et al., 2017), we showed that the response latency through the GFS (i.e. the time between the stimulation of the GFs in the brain, and flight or jump muscle depolarization) increases with age, demonstrating an age-related decline in the functionality of the escape circuit. Our experimental results suggest that the prolonged signal propagation is likely due to the age-associated decline in the conductance via gap junctions. This hypothesis is based on our findings that the old flies exhibited severely reduced ShakB plaque size, with other potentially contributing factors to this decline such as neuromuscular function and GF diameter being unaffected by age (Augustin et al., 2017). In this study, we generated a realistic computational biophysical model of the GFS based on these findings and on previously reported properties of the circuit’s components. By exploring potential determinants of response latency, including membrane properties, neuronal geometry and gap junction conductance, we created a model that not only recapitulates our experimental results, but also elucidates the relative importance of different physiological and anatomical parameters in regulating the speed of signal propagation through this escape response circuit.

## METHODS

The model was implemented in the NEURON simulation environment with Python (Hines and Carnevale, 1997; Hines et al., 2009). Source code and examples are available for download at https://senselab.med.yale.edu/modeldb/enterCode.cshtmPmodeh245415 (modelDB).

### Model architecture

The model is comprised of four cells: The Giant Fiber (GF), the tergotrochanteral muscle motoneuron (TTMn), a peripherally synapsing interneuron (PSI) and a dorsal longitudinal flight muscle motoneuron (DLMn, Fig. 2A). Each neuron contains 1-3 cylindrical compartments with dimensions based on anatomical data (see model parameters below). All compartments are divided into 51 segments each, and the membrane potential in each segment is calculated as a function of time based on the cable equation and any fixed or time-varying membranal conductances it contains. The GF is modelled as a single active compartment that forms unidirectional electrical synapses onto the active process (axon) of the PSI and the medial dendrite of the TTMn. The TTMn contains two dendrites (medial and lateral) (Godenschwege et al., 2002a) and an active axon. The PSI contains a dendrite and an axon, which forms a chemical synapse onto the active process (axon) of the DLMn. The DLMn contains a tapering axon and a dendrite (Egger et al., 1997; King and Wyman, 1980).

**Figure 2.**
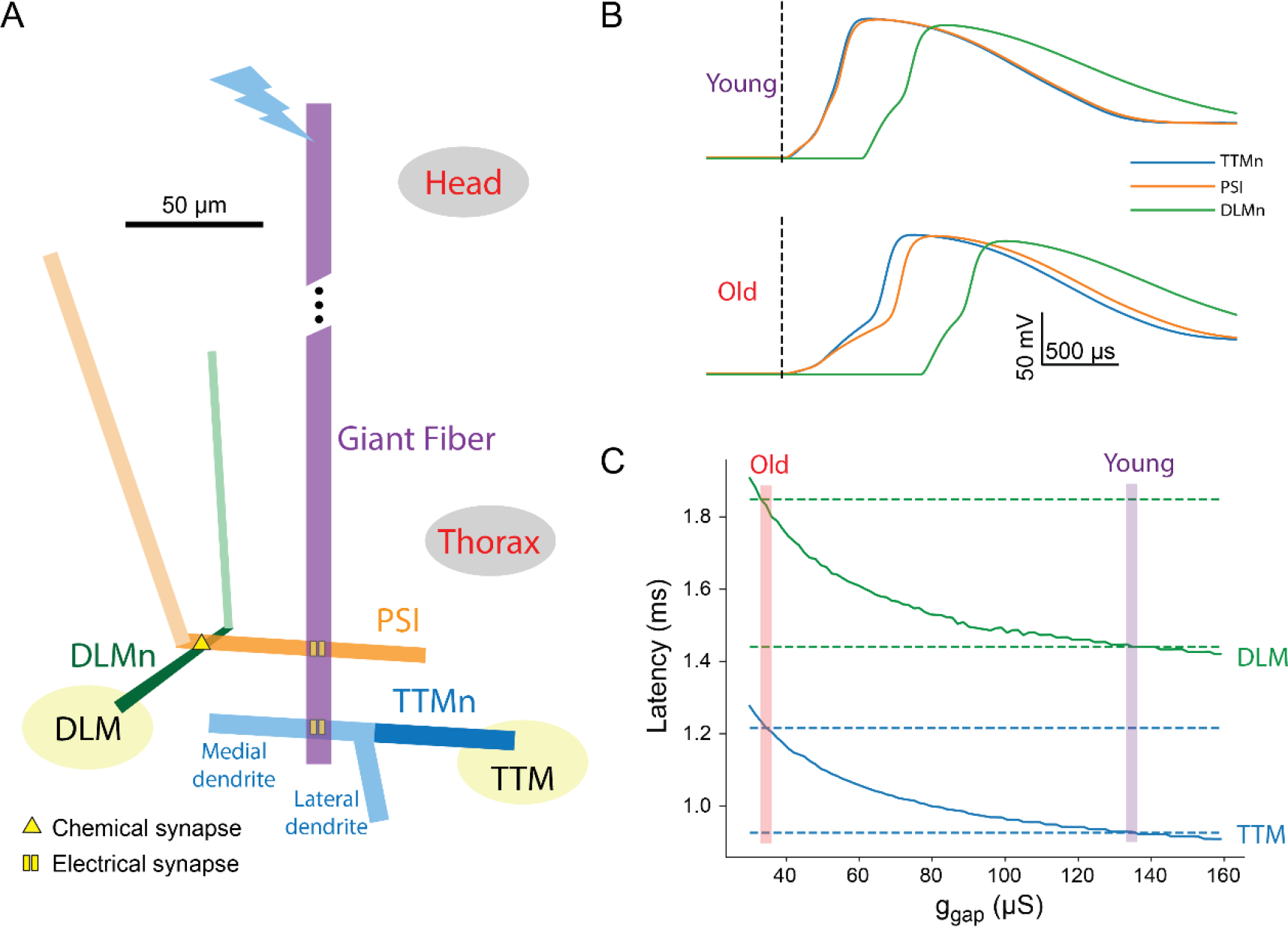
GFS model architecture and response latency measurements. **(A)** Model architecture and geometry, showing the cylindrical compartments that make up the four cell types in the model (to scale), along with the location of electrical and chemical synapses. Active compartments (axons) are shown in dark colours. Bolt denotes the proximal end of the GF that is stimulated in the simulation. The response latency in the DLM pathway is slightly delayed compared to the latency in the (shorter) TTM branch. **(B)** Membrane potential recorded in the model TTMn (blue), DLMn (green) and PSI (orange), for ‘young fly’ g_gap_ value (135 μS, top) and ‘old fly’ g_gap_ value (34.5 μS, bottom). **(C)** Latency from stimulus onset to muscle response as predicted by the model for TTM (blue) and DLM (green), as a function of g_gap_. The latency values recorded experimentally are indicated by dashed lines, and the g_gap_ values where they coincide with the values predicted by the model are shown by magenta and red bars (for young and old flies, respectively).

### Model conductances

All model compartments contain a passive leak conductance. Active processes (axons) were largely modelled according to an existing model of a *Drosophila* motoneuron (Gunay et al., 2015) and based on Hodgkin-Huxley type channel kinetics (Hodgkin and Huxley, 1952). They contain persistent and transient voltage-gated sodium channels, as well as voltage gated potassium channels, with kinetics based on Günay et al., 2015. Each conductance type is distributed with equal density in all active processes. The PSI-DLMn chemical synapse is modelled as a double-exponential process.

### Simulation

To test the TTM and DLM response latency in the model, we stimulated the proximal end of the GF with a current step duration of 0.03 ms (similar to that used by Augustin et al., 2017) and amplitude of 120 nA to approximate the input to the GF during high amplitude head stimulation, and measured the latency to the action potential peak in the TTMn and DLMn. To compare this latency with the latency values measured experimentally, we added 0.35 ms to this value to account for the neuromuscular junction delay. This value was estimated from the experimentally measured “neuromuscular latency” of ^~^0.65 ms. The “neuromuscular latency” is the time period between thoracic stimulation that directly stimulates the motoneurons, and TTM or DLM (muscle) depolarization (Augustin et al., 2017). This duration is composed of the time that passes from stimulus onset to action potential peak in the TTMn or DLMn, and the time from the peak to muscle depolarization (NMJ delay). To estimate the first part, the model TTMn was stimulated directly, resulting in ^~^0.3 ms from stimulus onset to action potential peak; the estimated NMJ delay is the remaining 0.35 ms. This delay contributes a fixed bias to the latency values, and therefore plays no role in assessing the relative importance of model parameters.

### Model parameters

The model parameters were chosen according to known values from the literature, where available (Table 1). Some of these values were manually adjusted to make sure all the model cells are spiking, and the response latencies match recorded values. Dimensions in the simplified anatomy were chosen to capture the general proportions of the cells and the ratio between active and passive membrane area.

**Table 1.** Anatomical and physiological parameters used in the paper.

| **Anatomical parameters** |  |
| --- | --- |
| GF diameter | 8 $\mu\text{m}$ (Augustin et al., 2017) |
| GF length | 400 $\mu\text{m}$ (Phelan et al., 1996; Smith et al., 1996) |
| Distance of contact with TTMn from proximal end | 400 $\mu\text{m}$ (distal end of the GF) |
| Distance of contact with PSI from proximal end | 360 $\mu\text{m}$ (King and Wyman, 1980; Phelan et al., 1996) |
| TTMn diameter | 6 $\mu\text{m}$ (King and Wyman, 1980) |
| TTMn axon length | 50 $\mu\text{m}$ (Godenschwege et al., 2002b) |
| TTMn medial dendrite length | 60 $\mu\text{m}$ (Godenschwege et al., 2002b) |
| TTMn lateral dendrite length | 30 $\mu\text{m}$ (Godenschwege et al., 2002b) |
| Distance of input from GF from medial dendrite proximal end | 12 $\mu\text{m}$ (Godenschwege et al., 2002a; Godenschwege et al., 2002b) |
| PSI diameter | 4.5 $\mu\text{m}$ (King and Wyman, 1980) |
| PSI axon length | 90 $\mu\text{m}$ (Phelan et al., 1996; Egger et al., 1997) |
| PSI dendrite length | 170 $\mu\text{m}$ (estimated) |
| Distance of input from GF from PSI axon proximal end | 45 $\mu\text{m}$ (Egger et al., 1997, Blagburn et al., 1999) |
| Distance of contact with DLMn from PSI axon proximal end | 76.5 $\mu\text{m}$ (Egger et al., 1997) |
| DLMn dendrite and axon proximal diameter | 2 $\mu\text{m}$ (King and Wyman, 1980) |
| DLMn axon distal diameter | 4 $\mu\text{m}$ (King and Wyman, 1980) |
| DLMn axon length | 50 $\mu\text{m}$ (Sun and Wyman, 1997) |
| DLMn dendrite length | 100 $\mu\text{m}$ (Sun and Wyman, 1997) |
| Distance of input from PSI from axon proximal end | 12.5 $\mu\text{m}$ (Egger et al., 1997) |
| **Physiological parameters** |  |
| Leak conductance | 0.03 $\text{mS}/\text{cm}^2$ (estimated) |
| Maximal transient voltage gated sodium conductance ($\bar{g}_{\text{Nat}}$) | 300 $\text{mS}/\text{cm}^2$ (Gunay et al., 2015) |
| Maximal persistent voltage gated sodium conductance ($\bar{g}_{\text{Nap}}$) | 0.11 $\text{mS}/\text{cm}^2$ (Gunay et al., 2015) |
| Maximal voltage gated potassium conductance ($\bar{g}_{\text{K}}$) | 10 $\text{mS}/\text{cm}^2$ (estimated) |
| Gap junctions conductance ($g_{\text{gap}}$, young fly) | 135 $\mu\text{S}$ (estimated) |
| Gap junctions conductance ($g_{\text{gap}}$, old fly) | 34.5 $\mu\text{S}$ (estimated) |
| Chemical synapse rise $\tau$ | 0.1 ms (standard value) |
| Chemical synapse decay $\tau$ | 1 ms (standard value) |
| Chemical synapse reversal potential | 0 (standard value) |
| Chemical synapse delay | 0.15 ms (estimated) |
| Chemical synapse peak conductance | 80 $\mu\text{S}$ (estimated) |
| Neuromuscular junction delay | 0.35 ms (see Methods) |
| Leak reversal potential | -85 mV (Gunay et al., 2015) |
| Sodium reversal potential | 65 mV (Gunay et al., 2015) |
| Potassium reversal potential | -74 mV (Gunay et al., 2015) |

## RESULTS

### A conductance-based model of the Drosophila GFS reproduces ageing-related latency increase

To examine how the electrical coupling in the fly’s GFS contributes to the transmission latency, we stimulated the model circuit with a 120 nA pulse at the proximal end of the GF (Fig. 2A) and recorded the voltage at the distal ends of the TTMn, PSI and DLMn neurons. Setting the gap junction conductance (g_gap_) of all model electrical synapses to 135 μS resulted in the voltage recordings shown in Figure 2B (top). The latency from stimulus onset to action potential (AP) peak, summed up with a fixed neuromuscular junction latency (0.35 ms), matches the values recorded experimentally in the TTM and DLM of young (5-7 days old) flies (0.93 and 1.44 ms, respectively (Augustin et al., 2017). This value of g_gap_ will therefore be used to model the response latency in young flies.

Decreasing g_gap_ to 34.5 μS results in longer membrane charging time to firing threshold due to weaker current across the gap junction, and thus to increased latency for both TTMn and DLMn, up to the values recorded in old (45-50 days old) flies (1.22 and 1.85 ms, Fig. 2B, bottom). This value of g_gap_ will therefore be used to model the response latency in old flies. Scanning a range of conductance values reveals the expected monotonic decrease of the response latency as g_gap_ increases (Fig. 2C). The model therefore reproduces the time latencies from stimulus to jump and flight muscle depolarization and shows that a four-fold reduction in gap junction conductance by itself could account for the transition from the latencies measured in young flies to the latencies measured in old flies.

### Co-dependency of the response latency on g_gap_ and other physiological and anatomical parameters

Next, we tested how the response latency predicted by the model changes as a function of both g_gap_ and membranal conductance densities. We performed two-dimensional grid scans by varying g_gap_ (spanning the values used for young and old flies), along with either the maximal transient voltage-gated sodium conductance for all axons (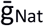, Fig. 3A), the maximal voltage-gated potassium conductance for all axons (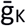, Fig. 3B), or the leak conductance in all processes (g_leak_, Fig. 3C). The latency values are presented as contour maps, where regions in the parameter space with a similar latency as young flies (± 1%) are denoted by red dashed lines. While the directionalities of the effects on response latency were expected from basic biophysical principles, this approach enabled us to assess the relative efficacies of these changes. As expected, increased transient voltage-gated sodium conductance reduced the response latencies (Fig. 3A) due to lowered AP threshold, and to a lesser degree by faster AP propagation. The reverse effect was observed when increasing the potassium and leak conductances (Fig. 3B-C), since these changes shift the membrane potential away from firing threshold and shunt inward current, leading to an increase in the time needed to reach threshold during a pre-synaptic spike. An extreme reduction in potassium conductance also prolonged the latency (Fig. 3B, left side of the plot) by elevating the resting membrane potential and thus causing sodium conductance inactivation. Within the tested ranges of conductance values, no change in a single parameter reverted the latency of an old fly (orange dot) to that of a young fly.

**Figure 3.**
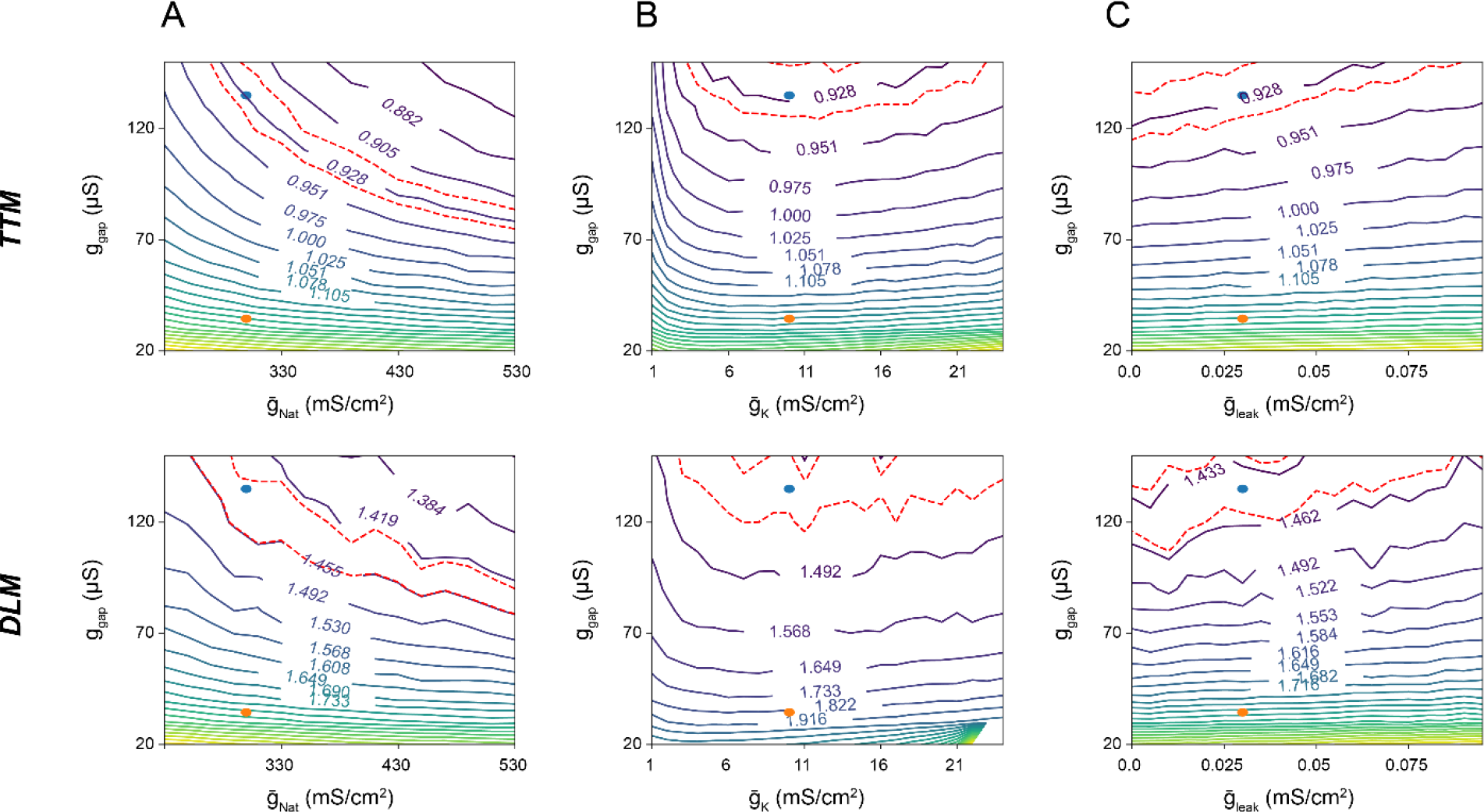
Co-dependency of the response latency on g_gap_. **(A-C)** The latency landscape, shown using iso-latency lines (labelled with response latency values in milliseconds) as a function of the global gap junction conductance (g_gap_) and maximal transient voltage-gated sodium conductance 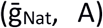, maximal voltage-gated potassium conductance 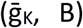 and leak conductance 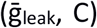, in TTM (top) and DLM (bottom). Blue and orange dots represent the values for young and old flies, respectively. The region in the landscape representing young fly latency is marked by red dashed lines.

We next tested how the response latency varied when changing g_gap_ along with anatomical parameters. For the TTMn branch of the circuit, we tested the following parameters: TTMn diameter (Fig. 4A), the length of lateral and medial TTMn dendrites (Fig. 4B-C), and the TTMn axon length (Fig. 4D). For the DLMn branch we tested the PSI diameter (Fig. 4E), PSI dendrite length (Fig. 4F), DLMn dendrite length (Fig. 4G) and the DLMn axon length (Fig. 4H). Since increases in diameter and length of neuronal processes decrease the cells’ input resistance, these changes in general decreased the effect of a given input current on the membrane potential and thus prolonged the latency to response. Among the tested parameters, a ^~^5-fold decrease in the PSI diameter reverted the response latency of an old fly to that of a young fly (Fig. 4E, black dashed line). Beyond a critical value, a further reduction in axonal length prolonged the response latency due to the decrease in active membrane surface area (and thus, in total active conductance, Fig. 4D and H). Overall, changes in the diameter of neuronal compartments affected the membrane capacitance throughout the length of the compartment, making their influence on the latency stronger compared to changes in compartment length. The results from Figures 3 and 4 show that, according to the model, changes in membrane conductance densities (within realistic limits) are far less efficient compared to anatomical changes in affecting the response latencies through the two branches of the GF circuit.

**Figure 4.**
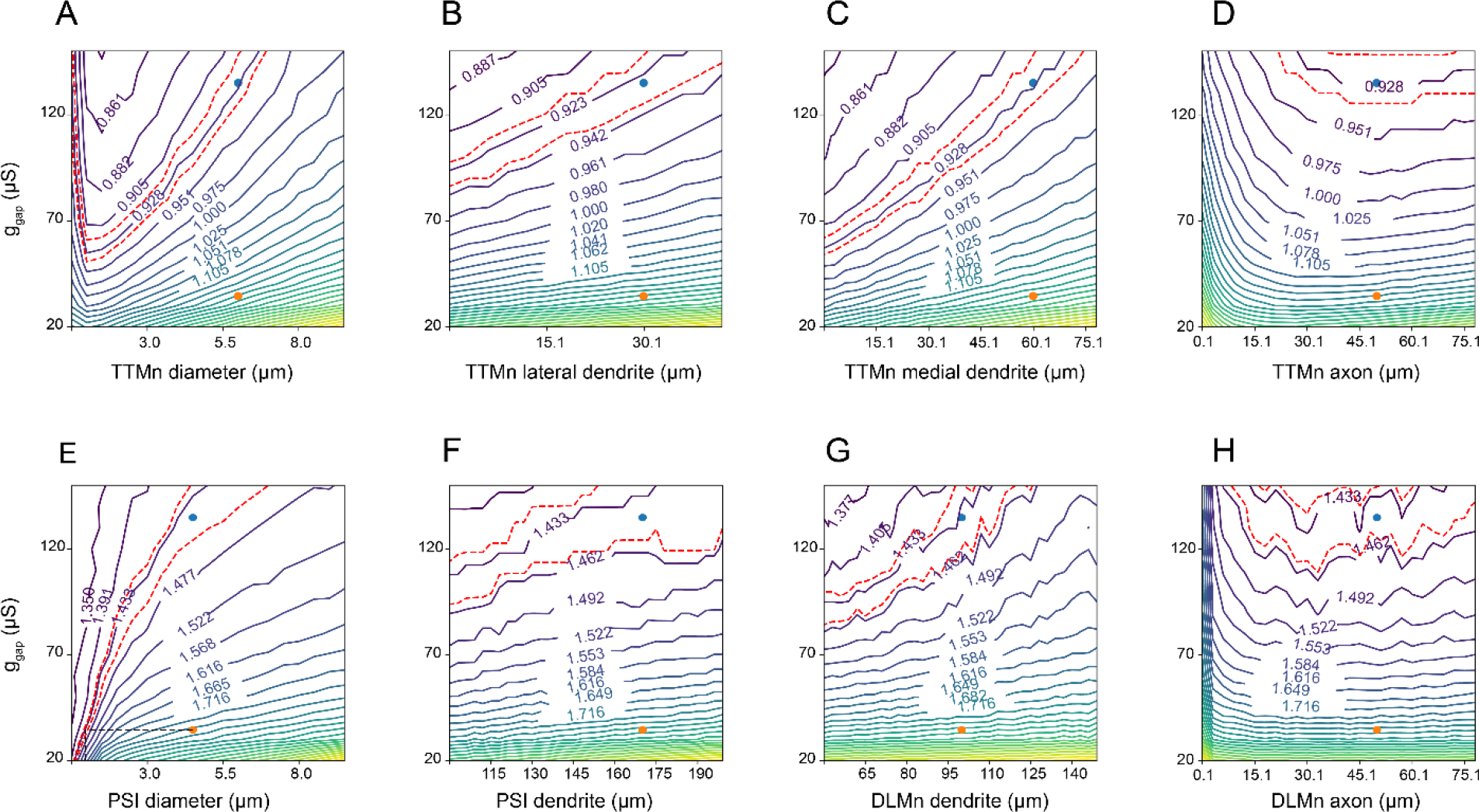
Impact of anatomical model parameters on response latency. **(A-D)** TTM Latency as a function of g_gap_ and anatomical parameters in the TTM branch of the model: the diameter of TTMn compartments (A), and the length of the TTMn lateral dendrite (B), medial dendrite (C) and axon (D). **(E-H)** DLM Latency as a function of g_gap_ and anatomical parameters in the DLM branch of the model: PSI compartment diameter (E), PSI dendrite length (F), DLMn dendrite length (G) and DLMn axon length (H).

### Co-dependency of the response latency on other parameter combinations

Our model enables predictions of the response latency as a function of arbitrary parameter combinations and may therefore be used as a benchside tool in experimental studies of signal propagation via the GFS. For example, Figure 5A shows the TTM latency landscape for voltage-gated sodium and potassium maximal conductances, each affecting the latency in a similar way as shown earlier against g_gap_ (Fig. 3A-B). Sodium and potassium reversal potentials, functions of intra- and extra-cellular ionic concentrations, influence the resting membrane potential, and thus their depolarization reduces the latency to TTM response (Fig. 5B). Increase in the TTMn medial and lateral dendrite lengths leads to an increase in membrane load and thus to increased TTM latency (Fig. 5C, see also Fig. 4A-H), but the medial dendrite length is more influential, since the GFs form synapses with the TTMn on this dendrite (Godenschwege et al., 2002a). The PSI-to-motoneuron contacts are chemical (Allen and Murphey, 2007; Tanouye and Wyman, 1980), so increasing the weight of this synapse naturally shortens the DLM response latency by accelerating the formation of action potential in the DLMn as a response to PSI activity; the synapse location on the DLMn dendrite has hardly any influence (Fig. 5D). We also tested the impact of the GF length and diameter on response latency through the GFS (Fig. 5E). For shorter axons (left side), the optimal diameter (where minimal latency is achieved) is rather small, since the longitudinal action potential propagation time is negligible compared to the passive charging of the membrane (see also Fig. 4A and E). For longer axons (right side), the propagation time becomes more important compared to membrane charging, so the optimal diameter is larger.

**Figure 5.**
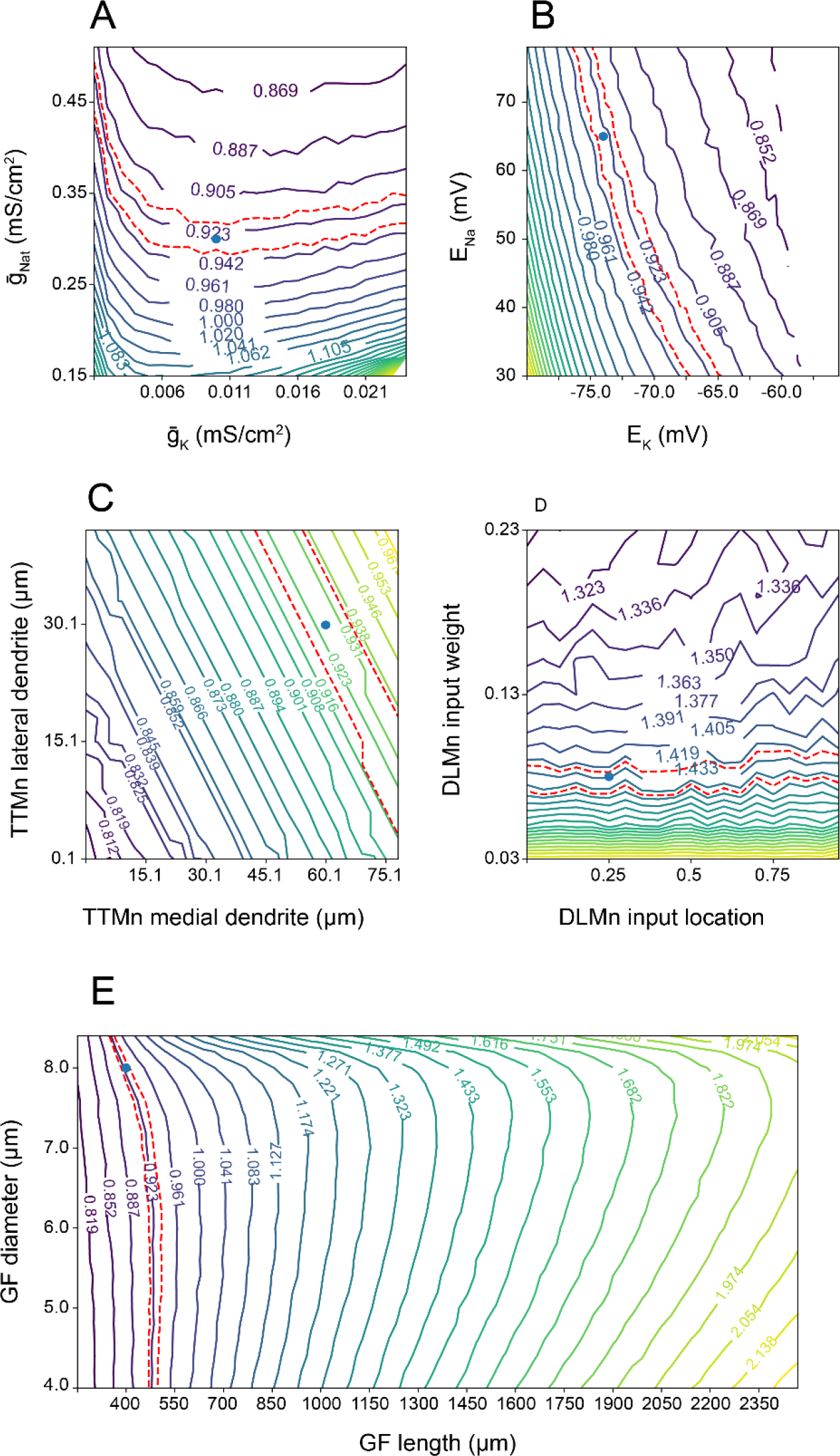
Co-dependency of the response latency on different parameter combinations. **(A)** TTM latency as a function of maximal voltage-gated transient sodium conductance 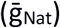 and maximal voltage-gated potassium conductance 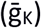. **(B)** TTM latency as a function of sodium reversal potential (E_Na_) and potassium reversal potential (E_K_). **(C)** TTM latency as a function of the TTMn medial dendrite length and TTM lateral dendrite length. **(D)** DLM latency as a function of the PSI-to-DLMn chemical synapse weight, and the synapse location along the DLMn dendrite. **(E)** GF latency as a function of the GF diameter and length. Blue dots represent the values for young and old flies, respectively.

In summary, among the tested parameter combinations we identified potassium reversal potential and dendrite length as the parameters with high impact on response latency, with the DLMn input location having a relatively small effect. These results can help guide experiments aimed at manipulating the GFS latency responses.

## DISCUSSION

Our biophysical model of the *Drosophila* GFS accomplished three main things. Firstly, it recapitulated the latency responses previously measured in young and old flies (Augustin et al., 2017). The so called “short-latency responses” are elicited by applying a high-frequency stimulus to the brain, thereby directly activating the Giant Fiber interneurons and bypassing the presynaptic (sensory) inputs to the GFs (Engel and Wu, 1996; Tanouye and Wyman, 1980). These “stimulus-to-muscle depolarization” response times therefore represent a readout for the functionality of the GFS that includes the GFs, the interneurons and motoneurons downstream of the GFs, as well as the (jump and flight) muscles innervated by the motoneurons (Fig 1). As the neuromuscular latency is not compromised by age (Augustin et al., 2017), we excluded the muscles (and their respective neuromuscular junctions) from the model, focusing on the period between the brain stimulus and motoneuronal (TTMn and DLMn) action potential peak. To be fully explained solely by the properties of gap junctions, our model suggests that the experimentally measured age-related increase in response latency requires a ^~^75% reduction in gap junctional conductance (Fig 2B-C). While decreased conductance can be caused by either gap junctional loss or dysfunction, this level of attenuation of conductance via gap junctions is within our estimate of the age-associated gap junctional loss in the GFS (Augustin et al., 2017).

Secondly, our model demonstrated the degree to which manipulations of principal membrane ionic conductances can influence the GFS response latencies. Augmentation of the transient voltage-gated sodium conductance in our model (equivalent to an overexpression or hyperactivation of voltage-gated sodium channels *in vivo*) shortened the response latency, with the increase in potassium and leak currents having the expected, opposite, effect (Fig. 3A-C and Fig. 5A). Considering the critical role of sodium currents in the generation of APs (Elmslie, 2010), the effect of increased Na^+^ conductance is likely due to lowered AP threshold. However, when manipulated within a physiologically relevant range, sodium conductances were unable to revert the latencies from old flies to youthful levels (Fig. 3A).

Thirdly, we showed that anatomical features of the GFS neurons can have a stronger effect on the speed of signal propagation via the circuit (Fig. 4 and Fig. 5C) compared to physiological parameters. Since the main factor in the model’s response latency is the membrane charging time to action potential threshold, any change to the membranal load (e.g. change in membrane surface area as a consequence of changing the length or diameter of a neuronal process) strongly affects the response latency by modulating the cells’ response dynamics to input current. Interestingly, decreasing the diameter of the PSI in old flies resulted in the reversal of response latencies to youthful values. For longer axons such as the Giant Fiber, however, the propagation velocity may be more important than the membrane charging time (which is influenced by the membranal load) (Rall, 2011). Indeed, optimal GF axon diameters scale with the length of axonal compartment in determining the GFS response latency (Fig. 5E).

Since the bulk of the signal propagation in the circuit is done via long axons and with minimal convergence of inputs, we could use simple, reduced geometry to represent the circuit components. With the addition of ion channel parameters previously characterized in *Drosophila* motoneuron simulations and several literature-based physiological constraints, our model is likely to faithfully reproduce salient anatomical and physiological features of the GFS.

The versatile *Drosophila* genetic toolbox allows for time-controlled modulation of relevant membrane conductances in individual GFS neurons and in the whole circuit (Jenett et al., 2012; Olsen and Wilson, 2008). Genetic manipulations of neuronal morphologies, including the size dendritic and axonal compartments, are also possible (Scott et al., 2003; Sugimura et al., 2004), although these interventions haven’t yet been tested in the GFS. These tools can be used to experimentally test the results presented here, with the goal of improving the circuit’s function and furthering our understanding of this escape response system. Even though our model is only partially constrained and some of its parameters had to be estimated, it is useful in developing intuition on which circuit elements are likely to have greater influence on the GFS response latency. The model code was made available and can be easily adapted to explore additional parameter combinations in addition to the ones presented here, and updated as more constraints become available.

## ACKNOWLEDGEMENTS

We would like to thank Konstantinos Lagogiannis (King’s College London) for providing useful comments on the manuscript, The Okinawa Institute of Science and Technology for organising the Computational Neuroscience course where this project was initiated, and Marcus Allen for the artwork in Figure 1.

## COMPETING INTERESTS

The authors declare no competing or financial interests.

## FUNDING

This work was funded by a Wellcome Trust Strategic Award to L.P. and by the Max Planck Society.

